# Multiple introgressions shape mitochondrial evolutionary history in *Drosophila paulistorum* and the *Drosophila willistoni* group

**DOI:** 10.1101/2020.09.17.301572

**Authors:** Guilherme C. Baião, Daniela I. Schneider, Wolfgang J. Miller, Lisa Klasson

**Affiliations:** Molecular Evolution, Department of Cell and Molecular Biology, Science for Life Laboratory, Uppsala University, Husargatan 3, 751 24, Uppsala, Sweden; Lab Genome Dynamics, Department Cell & Developmental Biology, Center for Anatomy and Cell Biology, Medical University of Vienna, Schwarzspanierstraße 17, 1090, Vienna, Austria

**Keywords:** mitochondria, *Drosophila*, evolution, genomics, introgression

## Abstract

Hybridization and the consequent introgression of genomic elements is an important source of genetic diversity for biological lineages. This is particularly evident in young clades in which hybrid incompatibilities are still incomplete and mixing between species is more likely to occur. *Drosophila paulistorum*, a representative of the Neotropical *Drosophila willistoni* subgroup, is a classic model of incipient speciation. The species is divided into six semispecies that show varying degrees of pre- and post-mating incompatibility with each other. In the present study, we investigate the mitochondrial evolutionary history of *D. paulistorum* and the willistoni subgroup. For that, we perform phylogenetic and comparative analyses of the complete mitochondrial genomes and draft nuclear assemblies of 25 *Drosophila* lines of the willistoni and saltans species groups. Our results show that the mitochondria of *D. paulistorum* are polyphyletic and form two non-sister clades that we name α and β. Identification and analyses of nuclear mitochondrial insertions further reveal that the willistoni subgroup has an α-like mitochondrial ancestor and indicate that both the α and β mitochondria of *D. paulistorum* were acquired through introgression from unknown fly lineages of the willistoni subgroup. We also uncover multiple mitochondrial introgressions across *D. paulistorum* semispecies and generate novel insight into the evolution of the species.

## 1. INTRODUCTION

Hybridization is increasingly recognized for its importance in promoting genetic diversity as well as for its consequent influence on adaptation and speciation. Large-scale genome sequencing is showing that the process is not only widespread but also relatively common, with many plants, fungi and animals carrying nuclear and cytoplasmic signatures of other species (Edelman and Mallet, 2021; Taylor and Larson, 2019). Genetic elements acquired through introgression — i.e., hybridization followed by back-crossing of hybrids to the parental lineages — often have adaptive value and lead to the emergence of novel traits or increased resistance to environmental stresses (Arnold and Kunte, 2017; Edelman and Mallet, 2021; Oziolor et al., 2019). Once introgressed, genetic elements persist in the receiving species through drift, selection or association with selfish elements that drive their spread (Dumas et al., 2013; Quilodrán, 2020; Seixas et al., 2018). The maternally inherited and reproduction-manipulating symbiont *Wolbachia*, for example, is known to cause sweeps of introgressed mitochondrial types (i.e., mitotypes) across host populations (Dumas et al., 2013; Jiggins, 2003; N Miyata et al., 2020).

The analysis of introgressed genetic elements can contribute to our understanding of past evolutionary events that shaped the present biodiversity (Ottenburghs, 2020). Sequence analysis often allows precise identification of hybridizing lineages and thus generates insight into the evolutionary history and biogeography of organisms of interest (Barlow et al., 2018; Meyer et al., 2012; Taylor and Larson, 2019). However, introgressed elements sometimes do not show strong similarity to any known organisms. “Ghost introgressions” derived from unsampled or potentially extinct species are relatively common and are often detected during the analysis of mitochondrial genomes (Ottenburghs, 2020; Toews and Brelsford, 2012; Zhang et al., 2019). Such introgressions, either from “ghost” or well-known species, commonly lead to deep mitochondrial divergence in clades with closely related nuclear genomes (Hirano et al., 2019; Horoiwa et al., 2021; Makhov et al., 2021; Zadra et al., 2021). If not properly identified, these introgressions can mislead evolutionary analyses (Ottenburghs, 2020; Toews and Brelsford, 2012). Mitochondrial genomic analyses can be further complicated by nuclear mitochondrial insertions (NUMTs) which are found in many organisms. NUMTs sometimes provide important phylogenetic information (Hazkani-Covo, 2009), but they may also create practical problems if mistaken for the true mitochondrial DNA (Calvignac et al., 2011). As in the case of undetected introgressions, unidentified NUMTs sometimes lead to phylogenies that do not represent the true relationship between lineages and to incorrect estimates of biological diversity (DeSalle and Goldstein, 2019; Kress et al., 2015).

*Drosophila* is a powerful model for studying introgression, as most lineages carry genomic elements acquired from other species or clades (Garrigan et al., 2012; Suvorov et al., 2022; Turissini and Matute, 2017). In some cases, up to 10% of the genome is estimated to have originated via introgression (Suvorov et al., 2022). Within *Drosophila*, young clades represent particularly interesting study systems, as the lineages which form them are more likely to hybridize or to have done so in the recent past (Coyne and Orr, 1989). One such clade is the *Drosophila willistoni* species group, a recent radiation of Neotropical fruit flies (Zanini et al., 2018) which comprises 24 species split into three subgroups: alagitans, bocainensis, and willistoni. (Bächli, 2019). Among these, the willistoni subgroup is the best studied and comprises several incipient species with varying degrees of inter- and intra-specific reproductive isolation (Burla et al., 1949; Ehrman and Powell, 1982; Winge, 1965). Due to this, the willistoni subgroup has become an important model for investigating reproductive isolation, speciation and hybridization (Burla et al., 1949; Civetta and Gaudreau, 2015; Dobzhansky and Pavlovsky, 1967; Ehrman and Powell, 1982; Mardiros et al., 2016; Perez-Salas and Ehrman, 1971; Schneider et al., 2019; Winge, 1965; Winge and Cordeiro, 1963).

One of the willistoni subgroup species, *Drosophila paulistorum*, is a species complex *in statu nascendi*. It comprises six semispecies — Amazonian (AM), Andean Brazilian (AB), Centro-American (CA), Interior (IN), Orinocan (OR) and Transitional (TR) — that show distinct but partially overlapping geographical distributions (Dobzhansky and Spassky, 1959). The semispecies express variable levels of reproductive isolation and exhibit both pre- and post-mating incompatibilities with each other (Dobzhansky and Spassky, 1959; Ehrman and Powell, 1982). Typically, females discriminate against males of other semispecies and occasional crosses do not produce offspring or result in hybrid infertility and male sterility (Dobzhansky et al., 1964; Ehrman, 1965; Miller et al., 2010). However, the TR semispecies and certain lines of other semispecies are more permissible and sporadically produce viable and, on rare occasions, fertile hybrids, at least under lab conditions (Dobzhansky et al., 1964; Malogolowkin, 1962). Thus, it is possible that gene flow between semispecies still exists or occurred in the recent past, when the semispecies were younger and barriers against gene flow were still incomplete (Perez-Salas et al., 1970). This complex evolutionary scenario has motivated several phylogenetic studies which produced a good understanding of the relationships between species of the willistoni subgroup (Gleason et al., 1998; Robe et al., 2010; Zanini et al., 2018). However, the relationship between *D. paulistorum* semispecies remains somewhat controversial, as nuclear and mitochondrial markers have often produced fairly different phylogenetic trees (Gleason et al., 1998; Robe et al., 2010; Zanini et al., 2018). These incongruences suggest that hybridization and mitochondrial introgression may have occurred between semispecies, but the small number and short sequence length of the markers available in previous studies have not allowed definite conclusions to be reached.

In the present study, we investigate the evolutionary history of mitochondria in *D. paulistorum*. We use complete mitochondrial genomes and draft whole-genome assemblies of 25 *Drosophila* lines belonging to six species of the willistoni group and three species of the closely related saltans group. By comparing mitochondrial and nuclear phylogenies we uncover a complex evolutionary history involving multiple mitochondrial introgressions, both ancient and recent. We discover that *D. paulistorum* mitochondria are polyphyletic and form two distinct non-sister clades, which we name α and β. With our analyses of the α and β genomes as well as of NUMTs found in the *D. paulistorum* genome, we show that the willistoni subgroup has an α-like mitochondrial ancestor. Interestingly, we also find evidence that both current *D. paulistorum* mitotypes are not native to the species and were likely acquired through introgression. The current α mitotype appears to have introgressed prior to the divergence of *D. paulistorum* and *D. equinoxialis*. As for the β mitotype, we suggest that it was acquired by *D. paulistorum* prior to semispecies diversification through a ghost introgression from an unsampled or extinct lineage of the willistoni group. Our results further reveal multiple introgressions across semispecies of *D. paulistorum*, generating new insight into the evolution of the species.

## 2. MATERIAL AND METHODS

### 2.1. Fly lines, DNA extraction and sequencing

*Drosophila* lines used in this study are summarized in **Table 1**. Flies were kept at 25±1°C on Formula 4-24 *Drosophila* instant food (Carolina, USA) with a 12-hour light-dark cycle. For each line, 20 ovaries were dissected from 3-day-old adult females and DNA extraction was performed using the Gentra Puregene Blood and Tissue kit (Qiagen). DNA samples used for PacBio sequencing were produced using the protocol described in (Ellegaard et al., 2013). Briefly, *Drosophila* flies were transferred to apple juice agar plates and allowed to oviposit for 2 hours, after which the eggs were collected, washed and dechorionated in 50% bleach and manually homogenized using a plastic pestle. Following centrifugation and filtration of the homogenate, the resulting cell pellet was subjected to whole genome amplification using the Repli-g midi kit (Qiagen) after which the DNA was purified using QIAamp DNA mini kit (Qiagen) according to the manufacturer’s recommendations.

**Table 1.**
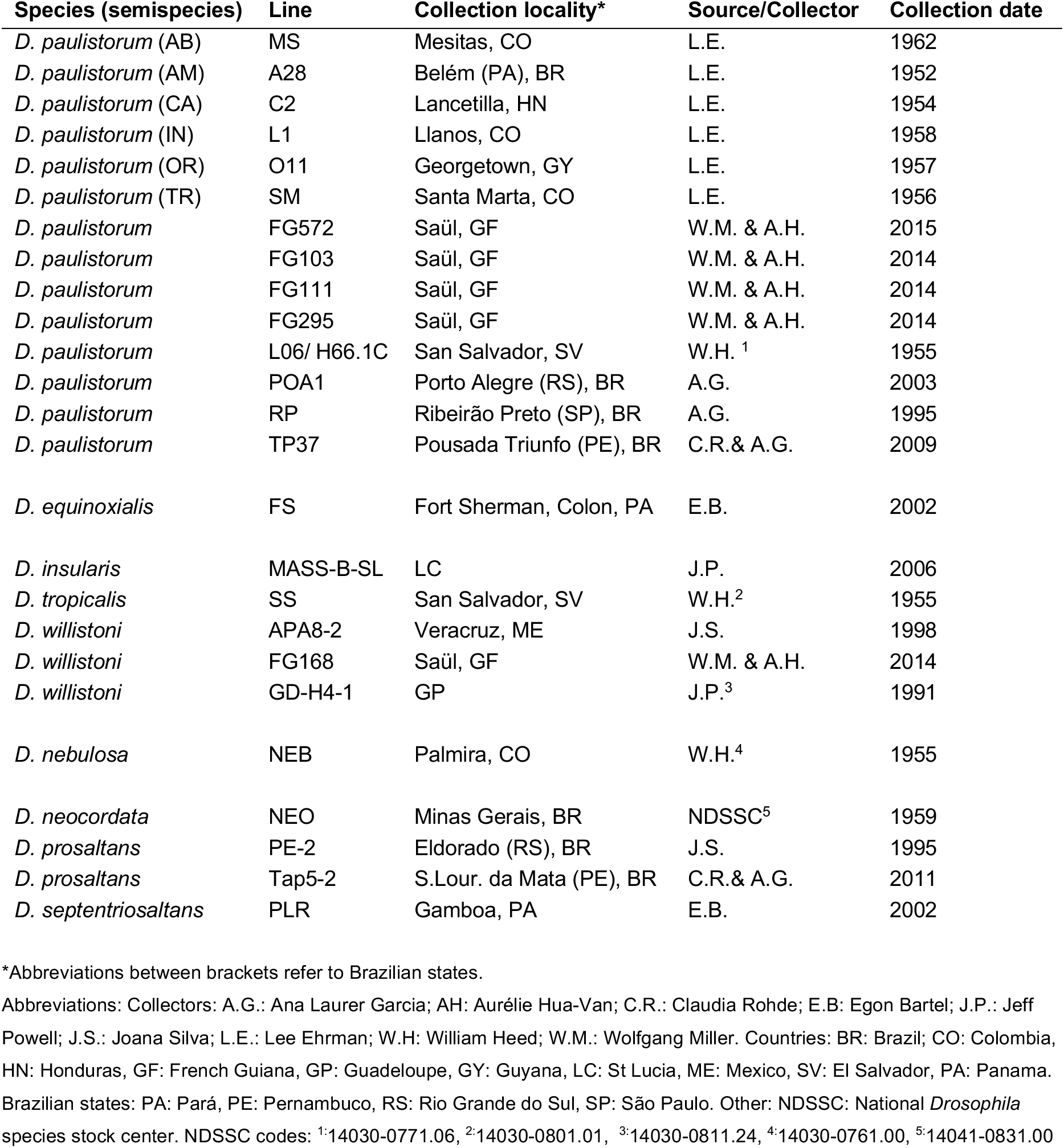
Fly lines used in this study.

#### 2.1.1. Sequencing

*Drosophila paulistorum* lines from the six classic reference semispecies plus lines from more recent collections were used for genome sequencing (**Table 1**). DNA extracted from ovaries was used to construct 350 bp fragment TruSeq libraries, multiplexed and sequenced at the Uppsala SNP and Seq platform in three different runs on an Illumina 2500 HiSeq machine generating 2×125 bp reads. The Illumina reads were quality filtered and trimmed using Trimmomatic-0.30 (Bolger et al., 2014) and error-corrected using SPAdes-3.5.0 (Bankevich et al., 2012). Amplified DNA extracted from early embryos of three lines (O11, POA-1 and TP37) was used to create 5 kb fragment SMRTbell libraries. Each library was run using P6-C4 chemistry in one SMRT cell on an RSII PacBio instrument. PacBio libraries were produced and sequencing was performed at the Uppsala Genome Center, Uppsala, Sweden.

### 2.2. Genome assemblies and annotation

#### 2.2.1. Mitochondrial genome assembly

The mitochondrial genome of the O11 line was generated by mapping all Illumina reads to the published mitochondrial genome of *D. willistoni* (NCBI accession number BK006338.1) using BWA-MEM v0.7.17 (Li, 2013) and thereafter extracting all aligned reads and their mates using SAMtools v1.14 (Li et al., 2009). The extracted Illumina reads were then subsampled to achieve an approximate coverage of 100 and assembled together with PacBio filtered subreads using SPAdes-3.10.1 with various k-mers settings. The genome came out as a single full-length contig. Both the Illumina and PacBio reads were then aligned to this contig using BWA-MEM v0.7.17. The resulting bam-file was converted to an ace-file that was used for manual curation of the sequence in Consed (Gordon et al., 1998). The mitochondrial genomes of all other *Drosophila* lines were generated by mapping Illumina reads to the O11 mitochondrial genome with BWA-MEM v0.7.17, extracting all aligned reads and their mates using SAMtools v1.14 and assembling the extracted reads with SPAdes-3.10.1 with various k-mers settings. Since PacBio data were only generated from the O11, TP37 and POA-1 lines, the assemblies of all other lines were run using only Illumina reads. For all lines except FG572, this procedure generated one contig that represented a complete or nearly complete circular mitochondrial genome. For FG572, this procedure created two nearly equal length contigs with similar coverage. The longest contig of ca. 14 kbp was similar to other β mt-genomes. The other large contig of ca 13 kbp was similar to α mt-genomes, and specifically to the NUMT that was discovered in the strain L06 (described in 3.3). Hence, this contig was regarded as a NUMT and not a real mitochondrion (see 2.6). To complete the mt-genome of FG572, we joined the 14 kbp β contig to a shorter contig of ca. 2000 bp which had almost double coverage and which represented the missing part of the mt-genome. The double coverage was due to the high similarity between the α-like NUMTand β mt-genome in this region.

In order to correct possible assembly errors, Illumina reads of each line were mapped back to the contig containing the complete or nearly complete mt-genome for that line using BWA-MEM v0.7.17 and indels were realigned with the IndelRealigner from GATK (McKenna et al., 2010). The resulting bam-file was then used to call variants using Freebayes with a range of parameters and converted to ace-format for manual curation in Consed (Gordon et al., 1998). In cases where high-frequency variants existed, we used the majority rule and base quality to select the consensus base.

The position and function of genes in the assembled mt-genomes were obtained by extracting the gene sequences from the published *D. willistoni* mt-genome (GD-H4-1 line) and using them as queries in a blastn search against each of our assembled mt-genomes. The start and stop positions of each gene were subsequently checked manually in Artemis (Rutherford et al., 2000) and adjusted if needed.

#### 2.2.2. Nuclear genome assemblies and retrieval of nuclear markers

Illumina reads from each *Drosophila* line were separately assembled with Megahit v.1.1.2 (Li et al., 2015) using multiple k-mer settings to generate draft nuclear genome assemblies. BUSCO v5.2.2 (Manni et al., 2021) was used for estimating the completeness of each assembly based on the ‘Diptera’ marker dataset. Complete single copy BUSCO markers which could be retrieved from all assemblies (**Table S3**) were then used for building nuclear phylogenies, as described in 2.3.

### 2.3. Alignments and phylogenetic analyses

Datasets of whole mitochondrial genomes, nuclear markers and mitochondrial genes were aligned with MAFFT v7.490 (mafft-linsi algorithm) (Katoh and Standley, 2013) and processed with TrimAl v1.4rev15 (Capella-Gutierrez et al., 2009) using the -automated1 flag for trimming poorly aligned regions. Concatenated sets of either mitochondrial or nuclear genes were created by combining trimmed alignments of single genes. Datasets of mitochondrial protein-coding genes containing either only the 3^rd^ or a combination of 1^st^ and 2^nd^ codon positions were created by parsing gene sequences using an in-house script. Alignments were visually inspected with Aliview v. 2019 (Larsson, 2014) for quality control.

Maximum likelihood trees were built with RAxML v8.2.12 (Stamatakis, 2014) using the flags: -f a -m GTRGAMMA -x 12345 -p 12345 -# 1000 (nucleotide trees) or: -f a -m PROTGAMMAGTR -x 12345 -p 12345 -# 1000 (amino acid trees). Bayesian trees were built with MrBayes v3.2.7a (Huelsenbeck and Ronquist, 2001; Ronquist and Huelsenbeck, 2003) using “nst= mixed” for estimating evolutionary models. Analyses were run for 10.000.000 generations with “nruns= 2”, “nchains= 4” and “samplefreq= 1000”. A burn-in of 25% was applied. Trees were visualized in FigTree v. 1.4.3 (Rambaut, 2009) and rooted using the species of the *saltans* group.

### 2.4. Quantification of synonymous and non-synonymous substitutions

In order to quantify synonymous and non-synonymous substitutions between the α and β clades, we first generated single gene alignments for all mitochondrial protein-coding genes using MAFFT v7.490. Synonymous and non-synonymous substitutions were counted in all possible pairwise comparisons between α– and β-carrying lines using codeml (PAML package v. 4.9j) (Yang, 2007). The values obtained were then averaged for each mitotype and gene. The same method was used for calculating the number of nonsynonymous substitutions per non-synonymous site (dN), synonymous substitutions per synonymous site (dS) and dN/dS ratios.

### 2.5. Recombination analysis

We checked for intergenic recombination between lines by analyzing incongruences between single gene trees which were built as described in 2.3. Tests for intragenic recombination on single-gene and whole-genome alignments were performed with Geneconv v.1.81 and PhiPack using default parameters.

### 2.6. Identification of nuclear mitochondrial DNA (NUMTs)

Using blastn, we screened our draft genome assemblies of willistoni subgroup lines for contigs which produced high-scoring pairs (HSPs) of at least 200 bp and 90% similarity to any of the assembled *D. paulistorum* mt-genomes. Among these, contigs with HSPs with 98% or higher identity to either α or β mt-genomes were tentatively assigned to those clades, while contigs with HSPs with 90-97% identity were deemed divergent. We further analyzed their similarity to the α and β mitotypes and calculated their coverage by using BWA-MEM v0.7.17 to map the Illumina reads of each line to their respective whole-genome assemblies combined with a representative of the α (C2) and β (O11) mitotypes. Read coverage was calculated with Mosdepth v0.3.2 (Pedersen and Quinlan, 2018) taking all mapped reads into account. Contigs with coverage lower than two were removed from the analysis to avoid potential artifacts. Contigs with similarity to mitochondria were considered NUMTs if their average coverage was similar (± 30%) to the nuclear coverage of their line. Additionally, contigs were classified as NUMTs if part of their sequence had similarity to the *Drosophila* nuclear genome, as indicated by blast searches against the NCBI nt database. The nuclear coverage of each line was defined as the average coverage of the 10 longest BUSCO markers used in our nuclear phylogenetic analysis (**Table S3**). We also calculated the percentage of bases of the α and β reference genomes that were covered by mapped reads using SAMtools mpileup. The result of these analyses is summarized in **Table S4**.

## 3. RESULTS

To investigate the evolutionary history of mitochondria in *D. paulistorum*, we sequenced and assembled the complete mitochondrial (mt) genomes and draft nuclear genomes of 23 *Drosophila* lines representing six species of the willistoni group and three species of the closely related saltans group (**Table 1**). Among these were 13 lines of *D. paulistorum*, including representatives of the six classic semispecies and seven recently collected isofemale lines from Brazil and French Guiana (**Table 1**). Additionally, we assembled the mt-genome of the *D. paulistorum* line L06/ H66.1C (**Table 1**), for which whole-genome Illumina data is publicly available (Kim et al., 2021).

The assembled mt-genomes are approximately 16.000 bp long, have a GC content of roughly 21% (**Table S1**) and encode the same 13 proteins, 22 tRNAs and two rRNAs as the mitochondria of *D. melanogaster* and most other animals. All mt-genomes have relatively short control regions (CR) of approximately 1.1 kb (willistoni group) to 1.3 kb (saltans group) compared to the 4.6 kb CR of *D. melanogaster*.

The draft nuclear assemblies have an average total length of 177 Mbp and vary considerably in the number of contigs, total length and quality (**Table S1**).

### 3.1. The phylogeny of *Drosophila paulistorum* and the willistoni group

We inferred a mitochondrial phylogeny of our fly lines based on our newly assembled mt-genomes and the mt-genome of the published *D. willistoni* reference genome GD-H4-1. The resulting tree revealed that *D. paulistorum* mitochondria are polyphyletic and split into two major clades, which we designate α and β (**Fig 1A**). The mt-genomes of the two clades are relatively distantly related and have an average nucleotide divergence of 2.4%. An analysis of substitution patterns between α and β showed that the rate of synonymous substitutions (dS) within protein-coding genes is higher than that of non-synonymous substitutions (dN) (**Table S2**), which indicates that purifying selection is acting on both clades.

**Fig 1.**
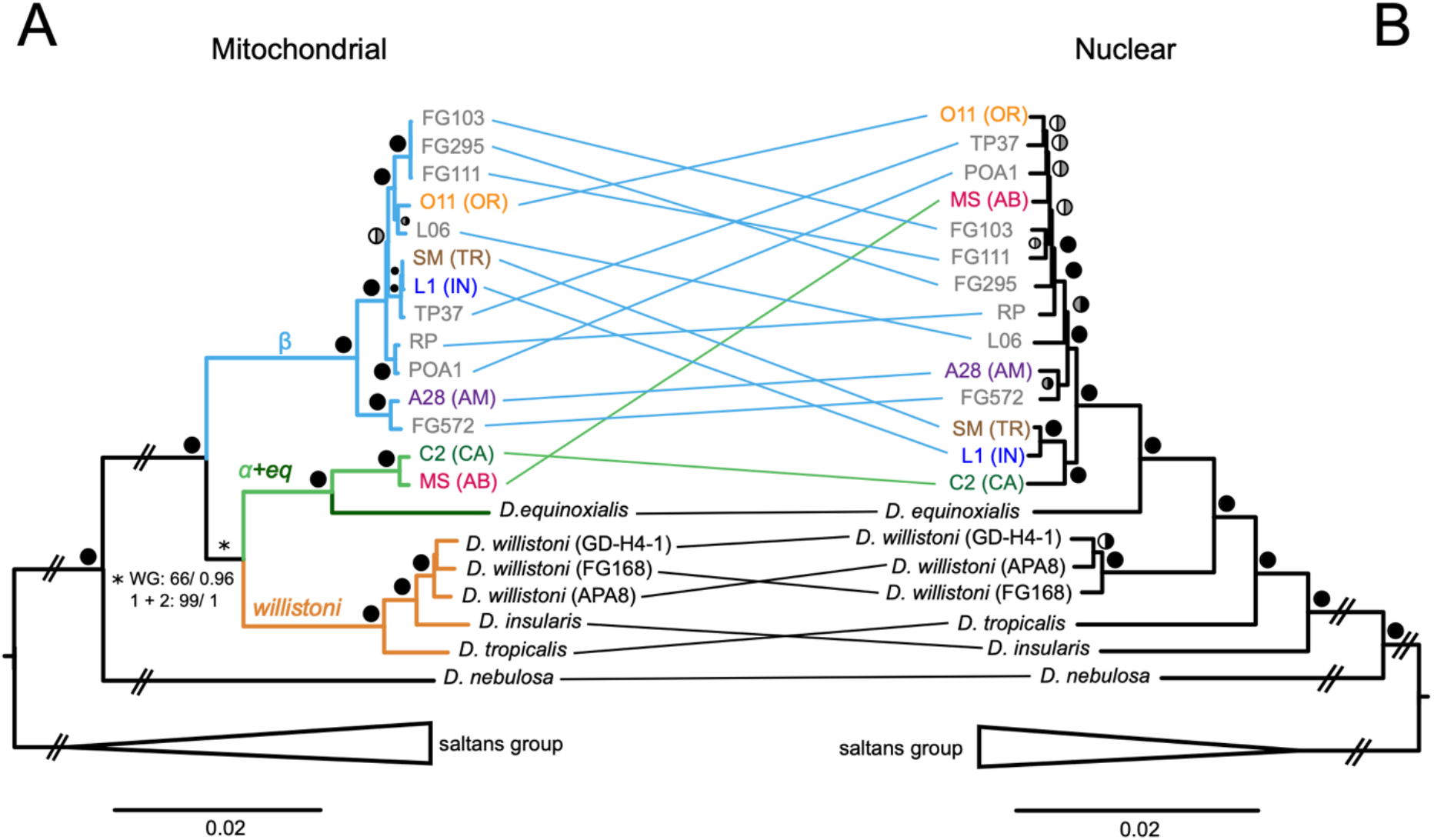
Phylogeny of *D. paulistorum* and the willistoni group. based on **(A)** whole mitochondrial genomes and **(B)** nuclear markers. *D. paulistorum* mitochondria form two non-sister clades: α and β. In the mitochondrial tree, the clade containing α and the mitochondria of *D. equinoxialis* (α+eq) is sister to the other species of the willistoni subgroup (willistoni clade). Support for this relationship is low in the whole genome (WG) phylogeny but high when amino acids or only the first and second codon positions of protein-coding genes are analyzed (1+2). The nuclear phylogeny shows *D. paulistorum* semispecies split into two sister clades with three semispecies each. For both trees, the same topology is obtained by maximum likelihood (ML, GTRGAMMA model) and Bayesian analyses. Colored circles next to the nodes indicate ML bootstrap support (left half) and Bayesian posterior probability (right half). Black represents 99-100% support, grey 70-98%, and white below 70%. Branches marked by transverse lines were shortened and do not follow the scale bar.

We found all deep nodes of the mitochondrial tree to be highly supported, except one (**Fig 1A**). The unsupported node is essential for our understanding of the phylogenetic relationships between *D. paulistorum* mitochondria since it defines the relationship between the clade which contains both the *D. paulistorum* α mitotype and the *D. equinoxialis* mt-genome (hereafter called “α+eq”) in regard to the clade containing the mt-genomes of *D. willistoni, D. insularis* and *D. tropicalis* (hereafter called the “willistoni clade”) (**Fig 1A**). Since further conclusions about the evolution of *D. paulistorum* mitochondria can only be made based on a robust phylogeny, we investigated possible underlying reasons for the low support of this node.

We tested if recombination between mitotypes could be affecting the phylogenetic signal by generating single gene trees for each of the 13 mitochondrial protein-coding genes and looking for conflicts between their topologies. We found that most of the trees had poor resolution due to the short sequence length of individual genes and low divergence between lines. Only two gene trees (ND2 and ND5) had good support for the node defining the relationship between α+eq and the willistoni clade (Bootstrap values > 80) (**Fig S1**). The ND2 tree featured α+eq as sister to the willistoni clade (BS = 80%) (**Fig S1A**) while the ND5 tree placed α+eq as sister to the β clade (BS = 92%) (**Fig S1B**). Among all single gene trees, *ND5* was the only one with support on all internal nodes, perhaps because it is the longest mitochondrial gene. We further tested for intragenic recombination in all protein-coding genes using two different algorithms and found that only *cytB* had a marginally significant signal (*p*<0.05) with one method, Phipack (see Methods). These weak and inconsistent signals indicated that recombination between mt-genomes is probably infrequent and thus unlikely to be the cause of the low bootstrap support in the tree.

Next, we tested if nucleotide composition bias could be responsible for the low bootstrap support. For that we built trees based on the concatenated sequences of the 13 mitochondrial protein-coding genes using either 1) amino acid (aa) sequences, 2) only the first and second codon positions or 3) only the third codon position. Support for α+eq as sister to the willistoni clade was strongly increased in the tree based on aa sequences (ML bootstrap 90%) in comparison to the whole mt-genome phylogeny (ML bootstrap 66%) (**Fig 1A, Fig S2A**). Using the first two codon positions of the nucleotide sequences of the protein-coding genes also resulted in considerably higher support for α+eq as sister to the willistoni clade (ML 99%, Bayesian 1) (**Fig 1A, Fig S2B**). In contrast, the tree based only on the third codon position showed high (Bayesian posterior probability 0.99) or fairly good (ML bootstrap 90%) support for α+eq and β being sister clades (**Fig 2**). Thus, given the conflicting topologies, we conclude that nucleotide composition bias is likely causing the low support of the node defining the relationship between the α+eq and willistoni clades in the mitochondrial phylogeny.

**Fig 2.**
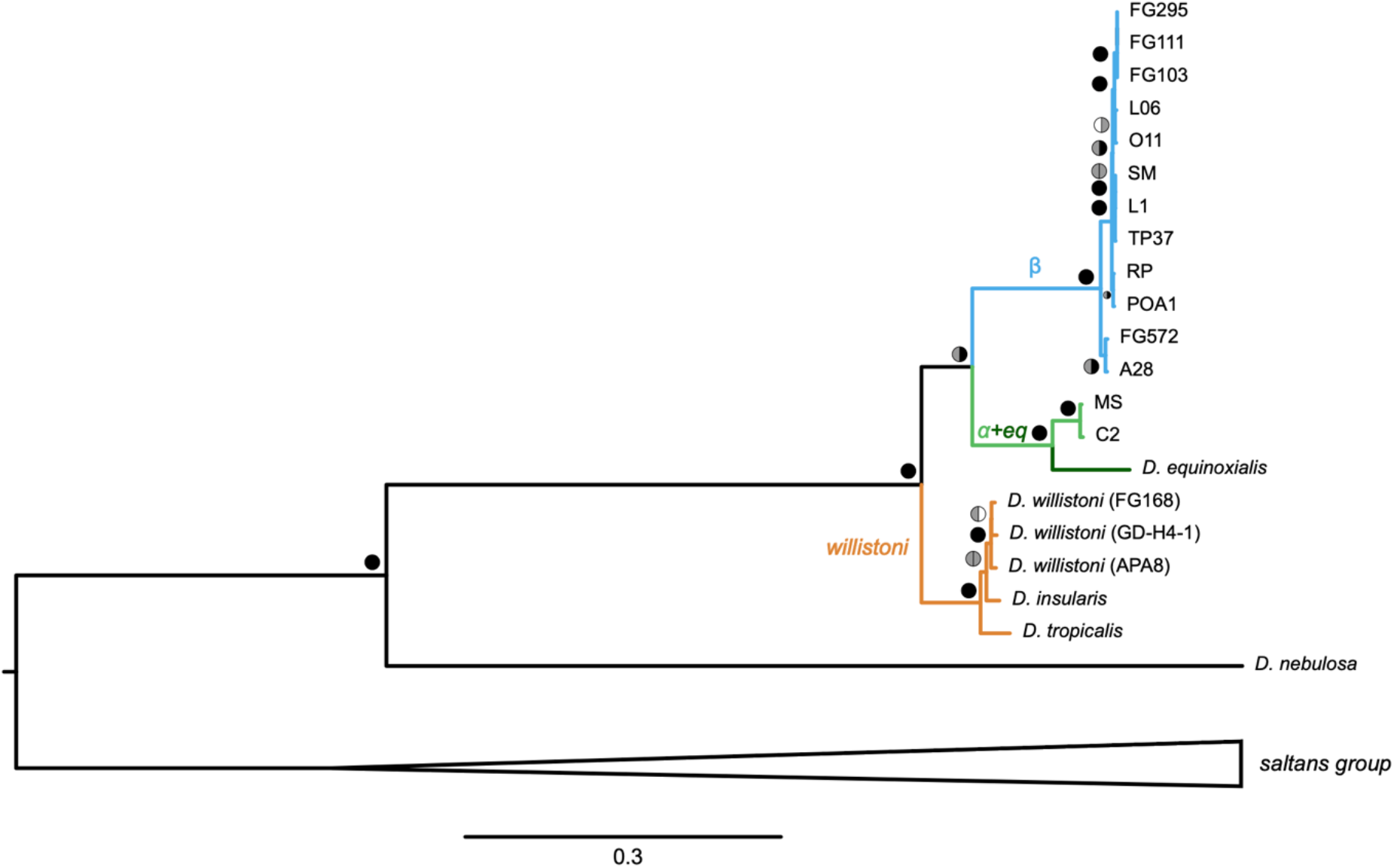
Mitochondrial phylogeny of *D. paulistorum* and the willistoni group based on the third codon position of the concatenated protein-coding gene sequences. The α+eq clade is highly supported as sister to β, which contrasts with its position as sister to the willistoni clade in trees based on whole mitochondrial genomes and on the first and second codon positions of the concatenated mitochondrial protein-coding genes (**Fig 1A, Fig S2B**). The change in topology is likely a result of convergence in nucleotide composition due to the ancestor of α+eq and β having lived in the same host. The same topology is observed in maximum likelihood (ML, GTRGAMMA model) and Bayesian analyses. Colored circles next to the nodes indicate ML bootstrap support (left half) and Bayesian posterior probability (right half). Black represents 99-100% support, grey 70-98%, and white below 70%.

Since the third codon position is more prone to bias than the first and second positions, we considered the topology supported by amino acids and by the first two codon positions to most likely represent the accurate evolutionary relationship between the clades. This is the same topology seen in the whole mt-genome tree (**Fig 1A**). Thus, we confirmed that the mitochondria of *D. paulistorum* are polyphyletic and that α+eq is sister to the willistoni clade rather than to β **(Fig 1A)**. The result also indicates that the ancestral mitochondrion of the willistoni subgroup is more similar to α (thus “α-like”) than to β.

To investigate the evolution of *D. paulistorum* mitochondria in more detail, we also inferred a reliable nuclear phylogeny that reflects the evolutionary history of the species and its semispecies (**Fig 1B**). The tree was based on amino acid sequences of 692 BUSCO markers (‘Diptera’ dataset) which were recovered as complete single copies in all of our draft nuclear assemblies (**Table S3**). The resulting nuclear phylogeny is highly supported in most non-terminal nodes and shows that the six classical semispecies of *D. paulistorum* are split into two sister clades, one formed by the AM, OR and AB semispecies and the other by the CA, IN and TR semispecies (**Fig 1B**).

### 3.2. Mitochondrial evolutionary history of *D. paulistorum* and the willistoni subgroup

A comparison of our nuclear and mitochondrial phylogenies (**Fig 1**) revealed inconsistencies that suggest both ancient and recent mitochondrial introgressions in *D. paulistorum*. First, the phylogenetic position of the β clade indicates that this mitotype was likely acquired by *D. paulistorum* from a donor species that diverged earlier than the willistoni subgroup but later than *D. nebulosa*. Second, the placement of the mitochondrial α+eq clade as sister to the willistoni clade is in strong contrast to the derived position that *D. paulistorum* and *D. equinoxialis* occupy within the willistoni subgroup in the nuclear phylogeny (**Fig 1**). This indicates that both extant mitotypes of *D. paulistorum* were potentially introgressed from species that branch outside of the willistoni subgroup.

Third, more recent mitochondrial introgressions appear to have occurred between the semispecies of *D. paulistorum*. This is evident by the fact that lines from different and relatively distantly related semispecies (based on our nuclear phylogeny) share the same mitotype, as is the case of the α mitotype in C2 (CA) and MS (AB) and the β mitotype in O11 (OR), L1 (IN) and SM (TR) (**Fig 1A**).

Finally, we note that *D. tropicalis* and *D. insularis* switch positions between the nuclear and mitochondrial phylogenies (**Fig 1**).

### 3.3. NUMTs provide further insight into mitochondrial evolution of the willistoni subgroup

While assembling the *D. paulistorum* genomes, we discovered that several of our lines likely carry nuclear mitochondrial insertions (NUMTs). We analyzed these to verify the accuracy of our assembled mt-genomes and to better understand mitochondrial evolution within the willistoni subgroup. Using our whole-genome assemblies, we identified contigs containing sequence fragments that showed at least 90% sequence similarity to representatives of the α and the β mitotypes of *D. paulistorum*. We then investigated their coverage and further analyzed their similarity to α and β by mapping reads of each line to their respective whole-genome assemblies plus representatives of α and β mt-genomes (see 2.6, **Table S4**). Contigs were classified as NUMTs if they either had a coverage similar to that of known nuclear genes or if they contained sequences identified as being of nuclear origin, regardless of coverage. The identified potential NUMTs exhibited varying lengths and degrees of sequence similarity to the assembled mt-genomes. Three of them were of particular interest for interpretations regarding mitochondrial evolution in *D. paulistorum*.

First, in several of the *D. paulistorum* lines, we identified a NUMT that lacks many of the typical hallmarks of a nuclear mitochondrial insertion. It spans the whole length of a mitochondrial genome (**Table S4**) and is almost identical to the α mitotype (99,4% nucleotide identity to a consensus of the C2 and MS mt-genomes). This atypical NUMT initially led us to believe that the lines which carry it were heteroplasmic. However, its identity as a NUMT was established due to a clear association with nuclear sequences in the published genome assembly of the *D. paulistorum* L06 line (Kim et al., 2021) as well as in contigs found in our draft whole-genome assemblies of the FG103, FG111, IN, O11 and SM lines. The high similarity between this NUMT and the α mitotype indicates that it is a recent nuclear transfer. However, most lines that carry it have a β mt-genome, suggesting that this NUMT does not originate from recent nuclear insertions of the resident mt-genomes. Thus, it must have been transferred between lines and semispecies in sporadic hybridization events. We are currently investigating the genomic and biological properties of this NUMT and intend to publish our results in a future publication.

The second NUMT of interest spans a genomic region that includes parts of the *ND2* and *CO1* genes. This NUMT was found in 13 of the 14 analyzed *D. paulistorum* lines and using it for phylogenetic analysis revealed a relationship between lines that is similar to that seen in the nuclear phylogeny (**Fig 3**, **Fig 1B**).

**Fig 3.**
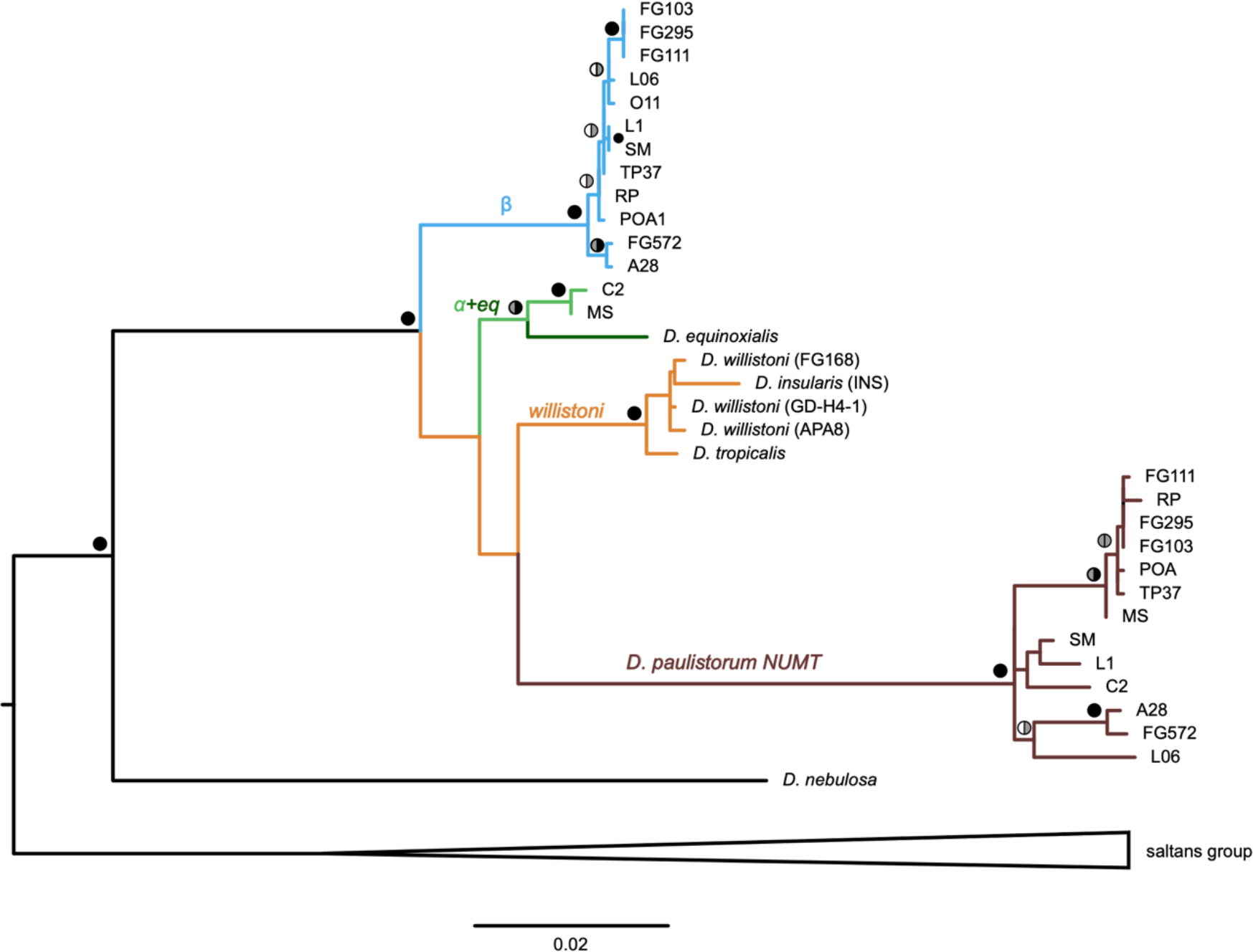
NUMT-derived phylogenetic relationship between *D. paulistorum* lines. The phylogeny is based on a NUMT sequence from *D. paulistorum* which covers part of the *ND2-CO1* genes combined with fragments of the true mt-genomes spanning the same region. The recovered phylogenetic relationship between *D. paulistorum* lines resembles that seen in the nuclear marker tree (**Fig 2B**), supporting an ancient origin of this nuclear insertion. The position of the NUMT next to the willistoni and the α+eq clades supports an α-like mitochondrial ancestor in the willistoni subgroup. Colored circles next to the nodes indicate ML bootstrap support (left half) and Bayesian posterior probability (right half). Black represents 99-100% support, grey 70-98%, and white below 70%.

This observation, combined with the presence of the NUMT in nearly all *D. paulistorum* lines, suggests that it stems from a single nuclear insertion that occurred before the diversification of *D. paulistorum* semispecies. Additionally, we noticed that the C2, L1 and SM lines share a unique insertion of a CR-1 family retrotransposon in this NUMT, further indicating that it is likely an old nuclear insertion. The presence of this retrotransposon also supports the monophyletic clade formed by the CA, IN and TR semispecies in our nuclear phylogeny (**Fig 1B**). The ancient origin of this NUMT combined with its phylogenetic position close to α and the willistoni clade (**Fig 3)** corroborates the hypothesis of an α-like ancestral mitotype in *D. paulistorum*.

Finally, we observed that the genome assembly of *D. equinoxialis* contains α few β-mitotype-related contigs which were not found in *D. insularis, D. tropicalis* or *D. willistoni*. The coverage of these contigs is similar to the nuclear average for the line but they do not contain matches to non-mitochondrial sequences. By investigating the coverage of these contigs, we saw that β-related reads present in our *D. equinoxialis* sample cover the whole extent of a mt-genome and result in a full mt-genome when assembled (**Table S4**). Based on the available data, we cannot be sure if these sequences stem from NUMTs or if they represent low-level contamination. Nevertheless, it is interesting to note that the presence of β sequences in *D. equinoxialis* might suggest, if true, that the last common ancestor of *D. paulistorum* and *D. equinoxialis* carried a β-like mitochondrion.

## 4. DISCUSSION

In this study, we investigate the evolutionary history of *D. paulistorum* using the complete mitochondrial genomes and draft nuclear assemblies of 25 *Drosophila* lines of the willistoni and saltans groups, including 14 *D. paulistorum* lines. We discover that *D. paulistorum* mitochondria are polyphyletic, forming two non-sister clades which we name α and β. We determine that the willistoni subgroup had an α-like mitochondrial ancestor and uncover multiple mitochondrial introgressions between *D. paulistorum* semispecies as well as across *D. paulistorum* and other unidentified lineages of the willistoni group. These corroborate previous findings of frequent hybridization in *Drosophila* (Suvorov et al., 2022) and add to reports of mitochondrial introgression across closely related *Drosophila*, as seen in the *D. yakuba* and *D. simulans* groups (Garrigan et al., 2012; Turissini and Matute, 2017). Using phylogenomics, we recover a nuclear phylogeny of *D. paulistorum* which is largely consistent with previous findings but does not recover either the AM or the TR semispecies as early-diverging and ancestral to the other semispecies, as proposed in other studies (Chao et al., 2010; Spassky et al., 1971; Zanini et al., 2018). Instead, we recover *D. paulistorum* semispecies split into two sister clades of three semispecies each. Our data also indicate that *D. paulistorum* carries several NUMTs, as previously seen in other *Drosophila* species including the relatively closely related *D. willistoni* (Rogers and Griffiths-Jones, 2012).

### 4.1. Evolutionary history of the α and β mitotypes of *D. paulistorum*

Our analysis shows *D. paulistorum* mitochondria to be polyphyletic and to form two non-sister clades that we name α and β. Both mitotypes show an early-diverging phylogenetic position which contrasts with the derived position of *D. paulistorum* within the willistoni subgroup (Gleason et al., 1998; Robe et al., 2010; Zanini et al., 2018). Thus, if our tree represents the true evolutionary history of these mitochondria, we can conclude that α and β are not originally from *D. paulistorum* but instead were introgressed from unknown early-diverging lineages. The same is true for the mitochondrion of *D. equinoxialis*, which is sister to the α mitotype (forming the α+eq clade) but branches outside *D. paulistorum* in the nuclear phylogeny. The positions of α+eq and of β in the mitochondrial tree suggest that both clades originate from lineages phylogenetically positioned between *D. nebulosa* and the willistoni subgroup, with the donor of β branching out earlier. We note that none of the sampled lineages of the willistoni group (except for *D. paulistorum*) has β-like mitochondria and that a search in GenBank reveals that α is the closest known relative to β. Hence, the donor lineage of β must be either still unsampled or extinct. One possibility is that α and β originate from rare and unsampled lineages in the alagitans or bocainensis subgroups of the willistoni group (Wheeler and Magalhães, 1962; Zanini et al., 2015). Most species in these subgroups are poorly known and have never been sequenced. Thus, it is possible that some occupy the likely phylogenetic position of the potential donors of α and β, i.e., independent lineages which are closer to the willistoni subgroup than to *D. nebulosa*. Ghost introgressions from unknown or extinct donors are reported in several organisms and often result in deep mitochondrial divergence among lineages with high nuclear similarity (Ottenburghs, 2020; Zhang et al., 2019), a situation comparable to what we observe in *D. paulistorum* and the willistoni subgroup.

Our analyses also allow us to infer at which point of the evolution of the willistoni subgroup the introgressions of α and β took place. Given the phylogenetic position of α as sister to the mitochondrion of *D. equinoxialis*, we conclude that the common ancestor of α+eq was introgressed into the common ancestor of *D. paulistorum* and *D. equinoxialis*. The potential β-NUMTs found in the genome of *D. equinoxialis* suggest that the same could be true for β. However, since we could not conclude if these β-like sequences in *D. equinoxialis* are indeed NUMTs or if they are the result of low-level contamination, it is also possible that β introgressed into *D. paulistorum* after the two species diverged. We note that β is present in 12 of the 14 sampled lines of *D. paulistorum* and four of its six semispecies. Thus, it is more parsimonious to assume that it was introgressed into *D. paulistorum* once, before semispecies diversification, rather than multiple times throughout evolution. Consequently, β must have been lost and replaced or failed to establish itself in the *D. paulistorum* lines which currently do not carry it i.e., C2 and MS. The introgression of β prior to *D. paulistorum* semispecies divergence is further supported by the similarity in nucleotide composition of the third codon position between the α and β mitotypes. Regardless of whether this convergence is caused by exposure of the β mt-genome to the mutational bias (amelioration) or the selective pressure (codon usage) of *D. paulistorum*, the effect is expected to be stronger when the genomes have been in the same host for a significant time (Marri and Golding, 2008).

One question that remains is which forces led to the successful establishment of the β mitotype in *D. paulistorum* and the replacement of its former mitotype. Since *D. paulistorum* originally had an α-like mitochondrion and considering that the genomes of α and β are relatively distinct from each other (2.4% divergence), it is likely that the introgression of β would have generated nuclear-mitochondrial incompatibilities (Burton et al., 2013). Such mitonuclear conflicts would in many cases lead to hybrid inviability, lower fitness or selection for the introgressed mitotype to be purged from the population (Burton et al., 2013). However, the β mitotype successfully established itself and spread in *D. paulistorum*. This suggests that the β mitotype either provided a strong fitness advantage in the conditions that prevailed at the time of the introgression or that its spread was driven by another factor, such as the endosymbiotic bacterium *Wolbachia* (Dean et al., 2003; Hill, 2019). *Wolbachia* strains that show strong reproductive manipulation or that increase host fitness sometimes spread fast through populations and may lead to hitch-hiking of mitotypes associated with infected females (Hurst and Jiggins, 2005; Turelli et al., 1992). In *D. paulistorum, Wolbachia* has been shown to affect fitness, mate choice and fecundity of some lines (Miller et al., 2010; Schneider et al., 2019). However, it is yet unknown if such effects occur in all semispecies or if they are directly associated with the spread of particular mitotypes in the species.

### 4.2. Mitochondrial introgressions between semispecies of *D. paulistorum*

Apart from the ancestral acquisition of the α and β mitotypes, *D. paulistorum* also shows signs of more recent mitochondrial introgressions across semispecies. These are indicated by incongruences between the mitochondrial and nuclear phylogenies. Given that *D. paulistorum* semispecies presently show pre- and post-mating incompatibilities with each other, it is likely that hybridization between them happened in an earlier evolutionary period when such barriers were less developed (Ehrman and Kernaghan, 1972; Ehrman and Powell, 1982). Potentially, sporadic hybridization may still occur in crosses involving the TR semispecies or populations of other semispecies which were shown to be more permissive under lab conditions (Dobzhansky and Pavlovsky, 1967; Ehrman, 1962; Malogolowkin et al., 1964). It is unknown if or how often *D. paulistorum* semispecies hybridize in the wild, but the fact that multiple semispecies can be found in sympatry shows that such events are not frequent enough to blur the distinctions between them (Perez-Salas et al., 1970).

We also observe some differences between our results and those of previous studies with regards to which mitotype is associated with certain semispecies, as inferred by an analysis of published phylogenetic trees. While here we see the IN semispecies carrying the β mitotype, it was previously associated with the α mitotype (Gleason et al., 1998) or with both α and β (Robe et al., 2010; Zanini et al., 2018) depending on the mitochondrial marker which was analyzed. Similarly, in this study, the TR semispecies carries the β mitotype, but it was previously associated with the α (Zanini et al., 2018) or with both the α and β mitotypes (Robe et al., 2010) depending on the marker used. Finally, while in our study the AB semispecies carries the α mitotype, it was associated with the β (Gleason et al., 1998; Zanini et al., 2018) or with both the α and β mitotypes (Robe et al., 2010) in previous studies. We note that the IN and TR lines used in the three mentioned studies were collected in the same sampling localities as the ones that we use in the present work, which suggests they may derive from the same lines. The same is true for the AB line used here and in Gleason et al. (1998) and Robe et al. (2010). However, the AB line used in Zanini et al. (2018) was collected in a different location — Santa Catarina, Brazil, while ours is from Mesitas, Colombia. Furthermore, all reference lines that were assayed in the three independent studies were obtained from different laboratories and at different time points. Thus, assuming that semispecies classifications are correct in all studies and that neither contamination nor permutation has occurred in any of the lines, we have two hypotheses to explain the conflicting mitotype associations. One possibility is that the incongruences are due to different populations or individuals from the same semispecies carrying distinct mitotypes i.e., some carry the α and some the β mitotype. Based on our nuclear phylogeny, we do not see evidence of intra-semispecies mitotype variation in our dataset. However, a wider sampling of lines and populations from each semispecies is required to conclude if such variation may occur. Alternatively, the differences between studies may derive from accidental amplification of NUMTs instead of actual mt-genome sequences. Amplification of NUMTs is a known issue when working with mitochondrial data and may misguide phylogenetic and barcoding analyses (Nacer and Raposo do Amaral, 2017; Song et al., 2008). Given this potential pitfall, we advise against using only mitochondrial markers for assigning *D. paulistorum* individuals and lines to semispecies. We also suggest that care should be taken when using solely mitochondrial markers to investigate poorly known lineages in general.

## 5. CONCLUSION

In the present study, we use whole genome sequencing combined with phylogenetics and comparative analyses to investigate the mitochondrial evolutionary history of *D. paulistorum* and the willistoni group. We discover that *D. paulistorum* mitochondria are polyphyletic, forming two distinct lineages, and find evidence that neither mitotype is original to the species. The multiple mitochondrial introgression events identified suggest that hybridization between *D. paulistorum* semispecies and across species of the willistoni group have occurred relatively frequently, both recently and in a more distant past. These conclusions are congruent with the recent evolutionary divergence of these lineages as well as their ongoing speciation and underline their value as models in hybridization and speciation studies. Our work also highlights the importance of genomic tools for elucidating past evolutionary and ecological events that contributed to forming the present biodiversity and its genetic variability.

## ACKNOWLEDGEMENTS

We thank Aurélie Hua-Van for fly sampling and the Nouragues research field station (managed by CNRS), which benefits from “Investissement d’Avenir” grants managed by Agence Nationale de la Recherche (AnaEE France ANR-11-INBS-0001; Labex CEBA ANR-10-LABX-25-01). Sequencing was performed at the SNP&SEQ Technology Platform and Uppsala Genome Center in Uppsala, Sweden, which is part of the Swedish National Genomics Infrastructure.

## DATA AVAILABILITY

Raw sequence data used in this project as well as assembled whole mitochondrial genomes are available in NCBI under the Bioproject PRJNA643793. Accession numbers of individual mitochondrial genomes are listed in **Table S1**.

## AUTHOR CONTRIBUTIONS

Guilherme Baião: Software, Validation, Formal analysis, Investigation, Data curation, Writing – original draft, Writing – review & editing, Visualization; Daniela Schneider: Investigation; Wolfgang Miller: Conceptualization, Resources, Funding acquisition, Writing - review & editing, Supervision, Project administration; Lisa Klasson: Conceptualization, Methodology, Validation, Formal analysis, Investigation, Resources, Data curation, Writing - original draft, Writing - review & editing, Visualization, Supervision, Project administration, Funding acquisition.

