## Supplemental figures for "Multiple introgressions shape mitochondrial evolutionary history in *Drosophila paulistorum* and the *Drosophila willistoni* group"

### **Supplementary Figures**

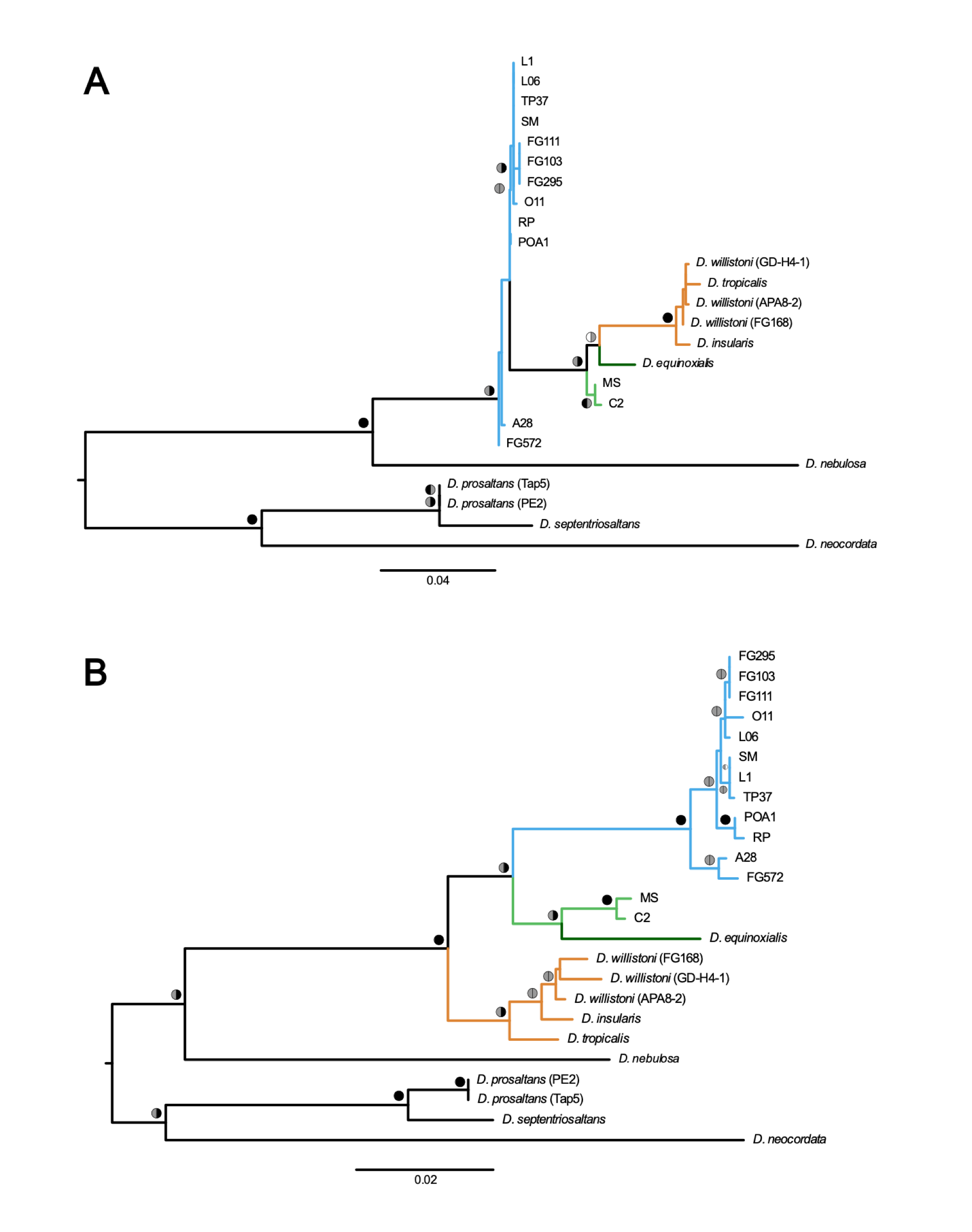

**Fig S1. Single-gene phylogenetic reconstructions** based on the (A) ND2 and (B) ND5 genes. These are the only mitochondrial genes whose single-gene phylogenies have good support on the node defining the relationship between α+eq and the willistoni clade. While the ND2 tree features α+eq as sister to the willistoni clade, ND5 shows α+eq as sister to the β clade. Colored circles next to the nodes indicate ML bootstrap support (left half) and Bayesian posterior probability (right half). Black represents 99-100% support, grey 70-98%, and white below 70%.

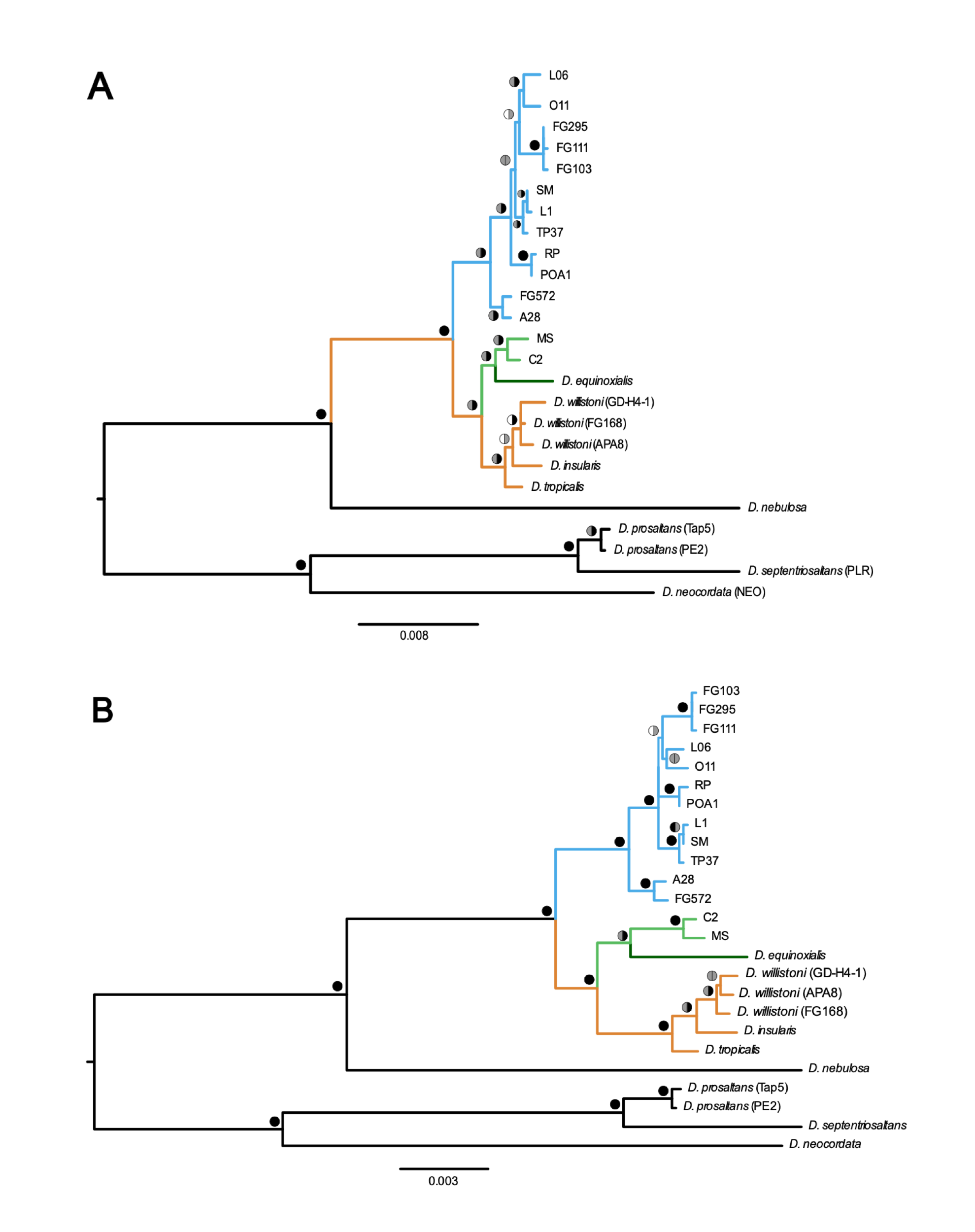

**Fig S2. Mitochondrial phylogeny of *D. paulistorum* and the willistoni subgroup based on concatenated (A) aminoacid sequences and (B) 1^st^ and 2^nd^ codon positions of the protein-coding genes.** Both trees show strong support for α+eq as sister to the willistoni clade. Colored circles next to the nodes indicate ML bootstrap support (left half) and Bayesian posterior probability (right half). Black represents 99-100% support, grey 70-98%, and white below 70%.
